# Brain Microstructure and Obesity Risk in Early Childhood: Insights from Restriction Spectrum Imaging

**DOI:** 10.1101/2025.09.19.677172

**Authors:** Mohammadreza Bayat, Bianca Braun, Madeline Curzon, Melissa Hernandez, Marilyn Curtis, Catherine Coccia, Dea Garic, Richard Watts, Mustafa Okan Irfanoglu, Paulo Graziano, Anthony Steven Dick

## Abstract

Pediatric obesity is a growing public health concern, yet little is known about the neurobiological underpinnings of obesity risk in early childhood. Using restriction spectrum imaging (RSI), we examined associations between adiposity and brain microstructure in a cross-sectional sample of 159 children aged 4–7 years, including 81 with ADHD and 78 typically developing (TD) peers. We focused on RSI-derived measures of restricted diffusion—specifically restricted normalized isotropic (RNI), directional (RND), and total (RNT) signals—as indicators of cellular density in subcortical and cortical regions implicated in reward and salience processing. Body mass index (BMI), percent body fat, waist circumference, and obesity status were assessed. Higher BMI, but not other adiposity measures, was significantly associated with increased RNI in the right insula, nucleus accumbens (NAcc), and putamen, as well as increased RNT in the right insula and pallidum. These findings suggest early microstructural alterations in reward-related circuits, consistent with theories of diet-related neuroinflammation. Contrary to hypotheses, ADHD diagnosis did not moderate the associations, and anthropometric profiles were similar between groups. This suggests a shared neural pathway linking early adiposity and brain structure, independent of diagnostic status. Our findings replicate and extend prior work in older children, highlighting BMI as the most sensitive marker of obesity-related brain differences in early childhood. These results underscore the potential of RSI as a tool for identifying early neural risk markers of obesity and inform future efforts to design preventive interventions during critical developmental windows.

## 1 INTRODUCTION

Pediatric obesity remains a growing public health concern, with rates rising steadily over the past several decades (Fryar et al. 2018). Obesity during childhood is linked to a range of adverse outcomes, including metabolic dysfunction, cardiovascular disease, and psychological distress (Reilly and Kelly 2011, Pulgarón 2013). Identifying early neurobiological predictors of obesity risk is critical for understanding underlying mechanisms and informing intervention efforts. Indeed, the nucleus accumbens (NAcc), a key region in the brain’s reward system, has been implicated in appetite regulation and susceptibility to weight gain (Volkow et al. 2013, Barbosa et al. 2022, Burger 2023). Recent studies by Rapuano et al. (2020, 2022) and Adise et al., (2025) found that increased cellular density in the NAcc, as measured via restriction spectrum imaging (RSI), predicted future weight gain in children. This marker is thought to reflect neuroinflammatory processes triggered by high-calorie food overconsumption, initiating a feed-forward cycle of excessive eating and metabolic disruption.

In addition to the NAcc, the insula has emerged as a region of interest in obesity research due to its role in taste perception and salience processing (Pritchard et al. 1999, Berthoud 2012, Rolls 2016). Neuroimaging studies show that the insula is reliably activated by palatable food cues, including in children and adults who are overweight or obese (Boutelle et al. 2015, Nakamura and Koike 2021, Avery et al. 2021, Doornweerd et al. 2018, Rapuano et al. 2023, Zhao et al. 2025). Structural differences in the insula have also been reported in pediatric samples, suggesting this region may contribute to early neural risk for obesity (Brady et al. 2025). However, the microstructural characteristics of the insula and their association with adiposity remain underexplored, particularly in young children.

Although Rapuano and Adise et al. (2020, 2025) focused on adolescents, it remains unknown whether there are similar associations between brain microstructure and adiposity earlier in development. Early childhood is a critical period for brain development and the formation of long-term eating behaviors (Dietz 1994, Liu et al. 2025), making it an ideal window for identifying neural risk markers. In addition, some evidence suggests children with attention-deficit/hyperactivity disorder (ADHD) exhibit higher obesity rates compared to typically developing (TD) children (Curzon et al. 2024, Cortese and Tessari 2017), although this effect may be small or negligible (Nigg et al. 2016, Turan et al. 2021). If present, such a pattern may reflect shared alterations in reward-processing circuits, as ADHD is characterized by dysfunction in mesolimbic dopamine pathways, including the NAcc (MacDonald et al. 2024, Nigg and Casey 2005, Volkow et al. 2009). However, it is unclear whether NAcc or insula microstructure differs between ADHD and TD groups, and whether these differences contribute to obesity risk.

To address these gaps, we conducted a cross-sectional replication study in a younger cohort (ages 4–7), including both TD children and those diagnosed with ADHD. Using RSI, we examined whether cellular density in the NAcc, insula, and other subcortical regions was associated with multiple anthropometric indicators: waist circumference, percent body fat, body mass index (BMI), and obesity status. Our aims were to (1) determine whether previously reported associations between NAcc microstructure and weight outcomes extend to early childhood, (2) assess whether these associations extend to insula, and (3) explore whether ADHD moderates these associations.

Based on prior findings, we hypothesized that increased NAcc and anterior insula cellular density would be associated with greater adiposity across all measures. We also hypothesized that these associations would be stronger in children with ADHD, consistent with theories linking reward system hyper-responsivity to impulsive eating behaviors (Yu et al. 2022, Barbosa et al. 2022).

This study contributes to the growing literature on the neural correlates of obesity risk in early childhood. By identifying brain-based markers that may predispose children to excessive weight gain, our findings could help inform early interventions aimed at modifying dietary habits before obesity becomes entrenched. Furthermore, by including children with ADHD, we offer a more nuanced view of how neurodevelopmental disorders may interact with reward and interoceptive systems to influence obesity risk. Together, this work extends prior findings to a younger, more developmentally sensitive population and highlights the potential for early brain markers—particularly in the NAcc and insula—to serve as indicators of emerging obesity risk.

## 2 METHOD AND ANALYSIS

### 2.1 Participants

The substudy was conducted at a university in the southeastern United States with a large Hispanic/Latinx population as part of a larger clinical intervention study for ADHD (Curzon et al. 2024). For the substudy, following data cleaning, 159 children had 1) usable diffusion-weighted and T1-weighted MRI scans (17 children were excluded due to excessive movement in the scanner; see Movement section), and 2) completed at least one anthropometric measure outside the MRI session. Of these, 81 children received a diagnosis of ADHD (see ADHD Diagnosis section). Some missing data were present due to the COVID-19 pandemic, which resulted in data loss for anthropometric measures requiring close personal contact. Missing data were addressed using multiple imputation (see Missing Data section).

#### Recruitment and Eligibility Requirements

Children and their caregivers were recruited from local schools, open houses/parent workshops, mental health agencies, and radio and newspaper ads. Exclusionary criteria for the children included intellectual disability (IQ lower than 70 on the Wechsler Preschool and Primary Scale of Intelligence 4th edition; WPPSI-IV (Wechsler et al. 2012)), confirmed history of autism spectrum disorder and/or Autism Spectrum Rating Scales Short Form score below clinical cutoff T-score of 60 (Goldstein and Naglieri 2009), and currently or previously taking psychotropic medication, including children who have been medicated for ADHD (i.e., all included children were medication naive). The study was reviewed and approved by the Florida International University Institutional Review Board.

#### Demographics of the Sample

Within this sub-sample, there were 78 typically-developing (TD) children and 81 children who were diagnosed with ADHD. ADHD diagnosis was accomplished through a combination of parent structured interview using the Computerized-Diagnostic Interview Schedule for Children (C-DISC (Shaffer et al. 2000)). The C-DISC is a highly structured, computerized diagnostic interview which can be used to assess over 30 different psychiatric disorders, including ADHD. It consists of a series of questions which, after a minimal training period, can be administered by “lay” interviewers, such as teachers and parents. In addition, we used parent and teacher ratings of symptoms and impairment using the Disruptive Behavior Disorders Rating Scale (DBD; (Fabiano et al. 2006a)), which involves teacher and parent reporting of symptoms of ADHD, including hyperactivity and impulsivity (Fabiano et al. 2006a), and the Impairment Rating Scales (Fabiano et al. 2006b). Children were classified as ADHD if raters endorsed clinically significant levels of ADHD symptoms (at least six symptoms of either Inattention or Hyperactivity/Impulsivity according to the Diagnostic and Statistical Manual of Mental Disorders Fifth Edition (DSM 5; (Association 2013)) or a previous diagnosis of ADHD), indicated that the child is currently displaying clinically significant academic, behavioral, and/or social impairments as measured by a score of at least 3 on a 7-point impairment rating scale. Dual Ph.D. level clinician review was used to verify diagnosis, supervised by P. G.. Thus, the final analyzed sample had the following demographics: (*M*_*age*_ = 5.46 years, *SD* = 0.69, 104 males (65%). The racial/ethnic breakdown of the subsample, which is based on United States National Institutes of Health demographic categories, was: 82% Hispanic/Latino; 87% White; 7% Black; 3% Asian; 3% More than one race. The distribution of maternal education was high school/less than high school = 3%; some college/Associate’s degree = 20%; college graduate = 35%; advanced degree = 41.5%.

### 2.2 Body Composition Measures

Four anthropometric measures were calculated: body mass index (BMI), percent body fat, waist circumference, and obesity status (dichotomous). Children height was measured (without shoes) to the nearest .01 cm using a wall-mounted stadiometer (Seca, Columbia, MD). The average of three height measurements was used for analyses. A Mediana i35 Body Composition Analyzer provided information regarding overall weight (to the nearest .01 kg) and body composition (i.e., percent body fat) via bioelectric impedance analysis. BMI was calculated using the height entered and the measured weight by dividing the weight in kilograms by the square of height in meters. BMI percentile and the BMI z score were calculated based on the age and sex norms of the Centers for Disease Control and Prevention (CDC) and the National Center for Health Statistics (Prevention 2024). Waist circumference was measured with a flexible tailor tape measure (in cm to the nearest 0.10 cm). Body composition variables (i.e., BMI and body fat percentage) were examined as continuous measures while overweight/obese BMI status was examined as a categorical outcome per CDC percentile cutoff scores (Prevention 2024).

### 2.3 Image Acquisition

All imaging was performed using a research-dedicated 3-T Siemens MAGNETOM Prisma MRI scanner (V11C) with a 32-channel coil located on the university campus. Children first completed a preparatory phase using a realistic mock scanner in the room across the hall from the magnet. Here, they were trained to stay still and were also acclimated to the enclosed space of the magnet, to the back projection visual presentation system and to the scanner noises (in this case, presented with headphones). When they were properly trained and acclimated, they were moved to the magnet. In the magnet, during the structural scans, children watched a child-friendly movie of their choice. Sound was presented through MRI-compatible headphones that also functioned as additional ear protection.

T1-weighted and DWI scans were acquired for each participant. T1-weighted MRI scans were collected using a 3D T1-weighted inversion prepared RF-spoiled gradient echo sequence (axial; TR/TE 2500/2.88; 1 x 1 x 1 mm, 7 min 12 s acquisition time) with prospective motion correction (Siemens vNav; (Tisdall et al. 2012)), according to the Adolescent Brain Cognitive Development (ABCD) protocol (Hagler Jr et al. 2019).

Movement artifacts pose challenges for T1-weighted images, which affects registration in the diffusion scan. Each T1 image was thoroughly reviewed by A.S.D. We applied a visual rating system ranging from ‘poor = 1’ to ‘excellent = 4’ for each T1-weighted image, with half-point allowances (e.g., 3.5). Most images were in the 3-4 range, with an average image rating of 3.54 (*SD* = 0.63). The ADHD and TD groups did not differ on image quality, *t*(144.7) = 1.11, *p* = 0.27.

DWI scans were acquired via high-angular-resolution diffusion imaging (HARDI) with multiband acceleration factor 3 (EPI acquisition; TR/TE = 4100/88 ms; 1.7 x 1.7 x 1.7 mm; 81 slices no gap; 96 diffusion directions plus 6 b = 0 s/mm^2^: b = 0 s/mm^2^ [6 dirs], b = 500 s/mm^2^ [6-dirs], 1000 s/mm^2^ [15-dirs], 2000 s/mm^2^ [15-dirs] and 3000 s/mm^2^ [60-dirs] s/mm^2^; A-to-P direction [7 m 31 s acquisition time]). A second brief scan at b = 0 s/mm^2^ in P-to-A direction was acquired to help deal with susceptibility artifacts.

#### Image Post-Processing

DWI preprocessing was performed using a number of complementary software suites. The steps were as follows: 1) DWI denoising (Veraart et al. 2016) and Gibbs ringing Correction (Kellner et al. 2016); 2) Inter-volume motion and eddy-currents distortion correction (Rohde et al. 2004, Pierpaoli et al. 2010) combined with volume-to-slice registration and outlier detection/replacement (Andersson et al. 2016 2017) as implemented in the TORTOISEV4 package (Pierpaoli et al. 2010); 3) creation of a synthesized T2-weighted image from the T1-weighted scan using Synb0-DisCo (Schilling et al. 2019 2020); 4) correction of spatial and intensity distortions caused by B0 field inhomogeneity, using TORTOISE DRBUDDI (Irfanoglu et al. 2015) implementing blip-up/blip-down distortion correction (Chang and Fitzpatrick 1992, Andersson et al. 2003, Morgan et al. 2004, Holland et al. 2010). This step uses both the reverse-phase encoded b = 0 s/mm^2^ image for the estimation of the field map, and the synthesized T2-weighted image for imposition of geometry constraints; 5) gradient non-linearity correction (gradwarp) using the gradient coefficients supplied by Siemens (Glover and Pelc 2019, Bammer et al. 2003, Barnett et al. 2021). The outlier-replaced, slice-to-volume registered, transformed for motion, eddy-current corrected, B_0_-induced susceptibility field corrected, gradient non-linearity corrected images are resampled to the T1-weighted resolution (to 1 mm_3_) and registered to the T1-weighted image in a single interpolation step. For whole-brain analysis, data are warped to the ABCD atlas space, and reported in LPS atlas coordinates (L = +; P = +; S = +), which are derived from the DICOM coordinate system. For ROI analysis, cortical surface and subcortical segmentation images of each T1 were reconstructed using Freesurfer (Destrieux et al. 2010, Fischl et al. 2004, Dale et al. 1999).

### 2.4 Movement

Participant movement has a substantial effect on DWI measurements, and can introduce spurious group differences in cases where none are present in the biological tissue, and it is recommended to incorporate movement as a nuisance regressor (Yendiki et al. 2014). To do this, we estimated movement using the root mean square (RMS) output from FSL eddy and implemented an overall movement cutoff for inclusion in analysis. This was arbitrarily defined as average RMS movement of more than one voxel (1.7 mm) over the course of the scan. In the subsample, 17 children (7.9%) were removed for excessive movement.

### 2.5 Restriction Spectrum Imaging

This study leveraged a multi-shell HARDI acquisition to implement Restriction Spectrum Imaging (RSI) (White et al. 2013a 2014 2013b, Brunsing et al. 2017), reconstructed using custom Matlab routines (Hagler Jr et al. 2019, Palmer et al. 2022). RSI allows for voxelwise estimation of the relative contributions of different diffusion components: restricted, hindered, and free water diffusion. These three compartments reflect the underlying microstructural environment. Free water, such as that found in cerebrospinal fluid, diffuses with minimal impedance. In contrast, biological tissue predominantly exhibits restricted and hindered diffusion (Bihan 1995). Hindered diffusion refers to water movement that is obstructed by cellular structures such as glial cells and neurites, typically following a Gaussian displacement profile. The signal arises from compartments larger than the diffusion length (about 10 *µ*m), influenced by how far water molecules must travel to circumvent structural obstacles. Restricted diffusion, in contrast, arises largely from cellular membranes and reflects molecular motion confined within tight cellular boundaries. Over longer diffusion times (Δ), the distinction between restricted and hindered diffusion becomes more pronounced (White et al. 2013a). The imaging parameters used here were selected to maximize sensitivity to these different types of diffusion behavior.

RSI enables the extraction of quantitative indices that reflect signal contributions from each diffusion compartment (Palmer et al. 2022). These indices are normalized to yield signal fractions, allowing for comparisons of restricted, hindered, and free water diffusion across voxels. In this study, we focused on four such metrics: the Restricted Normalized Directional signal (RND), the Restricted Normalized Isotropic signal (RNI), the Restricted Normalized Total signal (RNT), and the Hindered Normalized Total signal (HNT).

In the RSI model, restricted and hindered diffusion compartments are represented using fourth-order spherical harmonics, while free water diffusion is captured with zeroth-order harmonics. The axial diffusivity for restricted and hindered compartments is fixed at 1 × 10^-3^ mm^2^/s. Radial diffusivity is set to 0 mm^2^/s for restricted diffusion and to 0.9 × 10^-3^ mm^2^/s for hindered diffusion. Free water is modeled isotropically with a diffusivity of 3 × 10^-3^ mm^2^/s. Spherical deconvolution is used to estimate the fiber orientation distribution (FOD) in each voxel’s restricted compartment. RND is computed from the norm of the second and fourth-order harmonic coefficients, representing anisotropic diffusion across directions. RNI is derived from the spherical mean of the FOD, representing isotropic restricted diffusion. Together, RND and RNI sum to RNT. Further methodological details are available in previous publications (White et al. 2013a 2014 2013b, Brunsing et al. 2017, Palmer et al. 2022).

The RNT and HNT metrics, along with their components RNI and RND, serve as proxies for microstructural processes related to cellular and neurite development. These metrics reflect features such as myelination, dendritic arborization, and changes in neurite or cell body diameter or density, as well as increased concentrations of mature astrocytes. RND is thought to reflect anisotropic restricted diffusion associated with oriented structures like axons and dendrites, while RNI captures isotropic restricted diffusion, which may be linked to processes such as glial cell proliferation or changes in cell body structure. Prior research using similar acquisition parameters in children and adolescents (Palmer et al. 2022, Curtis et al. 2025) has demonstrated that RNT and HNT reliably distinguish between gray and white matter and show age-related variation in both tissue types. RNT and HNT tend to be inversely related—for example, increased myelination may elevate restricted diffusion (via reduced axonal membrane permeability) while decreasing hindered diffusion (due to reduced extracellular space). However, these relationships are modulated by tissue-specific characteristics and the distribution of diffusion signals across compartments within each voxel.

## 3 DATA ANALYSIS

### 3.1 Missing Data

In the final analyzed sample, a subset of anthropometric data was missing, including 13.8% of waist circumference values and 12.6% of percent body fat values. To address this, for the ROI analysis, we performed multiple imputation using fully conditional specification (FCS; (Van Buuren et al. 2006)) as implemented in the Best Linear IMPutation (BLIMP) software (Enders 2021). Twenty imputed data sets were defined. Convergence diagnostics indicated acceptable model performance, with potential scale reduction (PSR) values below 1.05 across chains, suggesting stable and replicable imputations. Parameter estimates were pooled according to Rubin’s rules (Rubin 1987, Li et al. 2025), using the mitml package in R (Grund et al. 2023).

### 3.2 Statistical Models

We conducted the statistical analysis using both regions of interest (ROI)-based analysis, and a voxelwise wholebrain analysis using a fast and efficient mixed-effects algorithm (FEMA; (Parekh et al. 2024)). Ordinary least squares linear models were specified for both ROI and voxelwise modeling. To be consistent across analyses, the diffusion metric (i.e., RNT, RNI, RND, HNT) was specified as the outcome, the anthropometric measure was specified as the main predictor, the diagnostic group was specified as the moderator, and additional covariates (age, sex, parent education, and movement) were added. If the moderating effect was not significant, we dropped the interaction term to simplify the model.

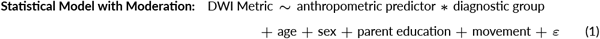

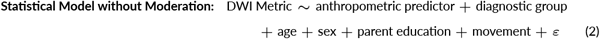

### 3.3 Regions of Interest Analysis

We identified the eight regions of interest from Rapuano and colleagues (2020) averaged across each hemisphere: NAcc, putamen, caudate, pallidum, ventral diencephalon, thalamus, amygdala, and hippocampus. We additionally added left and right anterior insula, based on our own review of the literature. We identified these regions anatomically on individual cortical surfaces using the automatic FreeSurfer parcellation (Destrieux et al. 2010, Fischl et al. 2004, Dale et al. 1999), based on the anatomical conventions of Duvernoy et al. (1999). These were corrected for multiple comparisons using False Discovery Rate correction (FDR; (Benjamini and Hochberg 1995)) across all ROIs within a measure.

For each region for each measure (RNI, RND, RNT, HNT) we applied the statistical model, adjusting the standard error estimates according to Rubin’s rules as described above. In each iteration, we checked whether the interaction effect was statistically significant after FDR correction (Model 1). If not, Model 2 was run dropping the interaction term.

### 3.4 Fast and Efficient Mixed-Effects Algorithm (FEMA) Analysis

We performed a supplemental voxelwise analysis using FEMA on the specified models (note, despite the name, ordinary least squares, not mixed-effects models, were run). The use of FEMA facilitates use of the ABCD atlas space, which maintains consistency with analyses by Rapuano and colleagues (2020). The design matrix is constructed identically to that of the ROI analysis. In addition, we masked on the whole-brain the ROIs applied on the ROI analysis. The per-voxel threshold was set at *p* < 0.01 (uncorrected) and is designed to show spatially where the most prominent results originate from the focused ROI analysis.

## 3.5 Results

We first investigated whether, in this sample, any of the anthropomorphic measures differed across TD and ADHD children. As Table 1 shows, although the direction of the effects indicated that ADHD children tended to score higher on all measures, none were statistically different, and the small effect sizes indicated no meaningful differences.

**TABLE 1.** Summary statistics for anthropometric measures.

| Measure | Group | M (SD) | *t* | *p* | Cohens *d* | CI for *d* |
| --- | --- | --- | --- | --- | --- | --- |
| BMI | ADHD | 16.68 (2.39) | -0.54 | 0.59 | -0.09 | [-0.40, 0.23] |
| BMI | TD | 16.48 (2.27) |  |  |  |  |
| % Body Fat | ADHD | 23.04 (8.11) | -0.61 | 0.54 | -0.10 | [-0.41, 0.22] |
| % Body Fat | TD | 22.31 (6.77) |  |  |  |  |
| Waist Circum. | ADHD | 57.93 (7.94) | -0.54 | 0.59 | -0.09 | [-0.40, 0.23] |
| Waist Circum. | TD | 57.34 (5.69) |  |  |  |  |

| Measure | Group | Proportion | *t* | *p* | $\phi$ | CI for $\phi$ |
| --- | --- | --- | --- | --- | --- | --- |
| Obesity Status | ADHD | 19.75% | 0.54 | 0.46 | 0.06 | [0.00, 0.21] |
| Obesity Status | TD | 14.10% |  |  |  |  |

Note. ADHD *n* = 81; TD *n* = 78.

We next investigated in all ROIs whether diagnostic group moderated associations between anthropomorphic measures and RSI measures (i.e., Model 1). None of the anthropomorphic by diagnostic group interaction effects survived the statistical correction. Thus, all reported ROI results are based on Model 2. Table 2 shows the results of this analysis that survived statistical correction. These effects were most reliable in right insula (for RNI, RND, and RNT), in NAcc (for RNI), and in putamen (for RNI) and pallidum (for RNT). Full uncorrected results are provided in Supplemental Tables 1-4.

**TABLE 2.** Associations of Body Mass Index with RSI Metrics that Survived Statistical Correction.

| RSI Predictor | Brain Region | $\beta$ | SE | *t* | *p* | 95% CI Lower | 95% CI Upper | *r<sub>sp</sub>* | FDR<sub>*p*</sub> |
| --- | --- | --- | --- | --- | --- | --- | --- | --- | --- |
| RNI | Right Insula | 0.22 | 0.08 | 2.80 | 0.01 | 0.07 | 0.38 | 0.22 | 0.03 |
| RNI | Nucleus Accumbens | 0.22 | 0.07 | 2.99 | 0.00 | 0.08 | 0.37 | 0.24 | 0.03 |
| RNI | Putamen | 0.18 | 0.07 | 2.68 | 0.01 | 0.05 | 0.31 | 0.21 | 0.03 |
| RND | Right Insula | 0.28 | 0.08 | 3.53 | 0.00 | 0.12 | 0.43 | 0.28 | 0.01 |
| RNT | Right Insula | 0.27 | 0.08 | 3.39 | 0.00 | 0.11 | 0.42 | 0.27 | 0.01 |
| RNT | Pallidum | 0.21 | 0.08 | 2.71 | 0.01 | 0.06 | 0.36 | 0.21 | 0.04 |

Note. All outcomes show results that survived FDR correction for the main effect of the brain predictor (with no interactions), controlling for diagnostic group, age, sex, parent education, and movement. The 95% CI is provided for $\beta$ . *r<sub>sp</sub>* is the semipartial *r*. FDR<sub>*p*</sub> is the False Discovery Rate corrected *p* value.

Figures 1 and 2 present these results visually, showing the strongest effects and associated confidence intervals. As shown, the strongest and most reliable effects were generally confined to the restricted diffusion measures.

**FIGURE 1.**
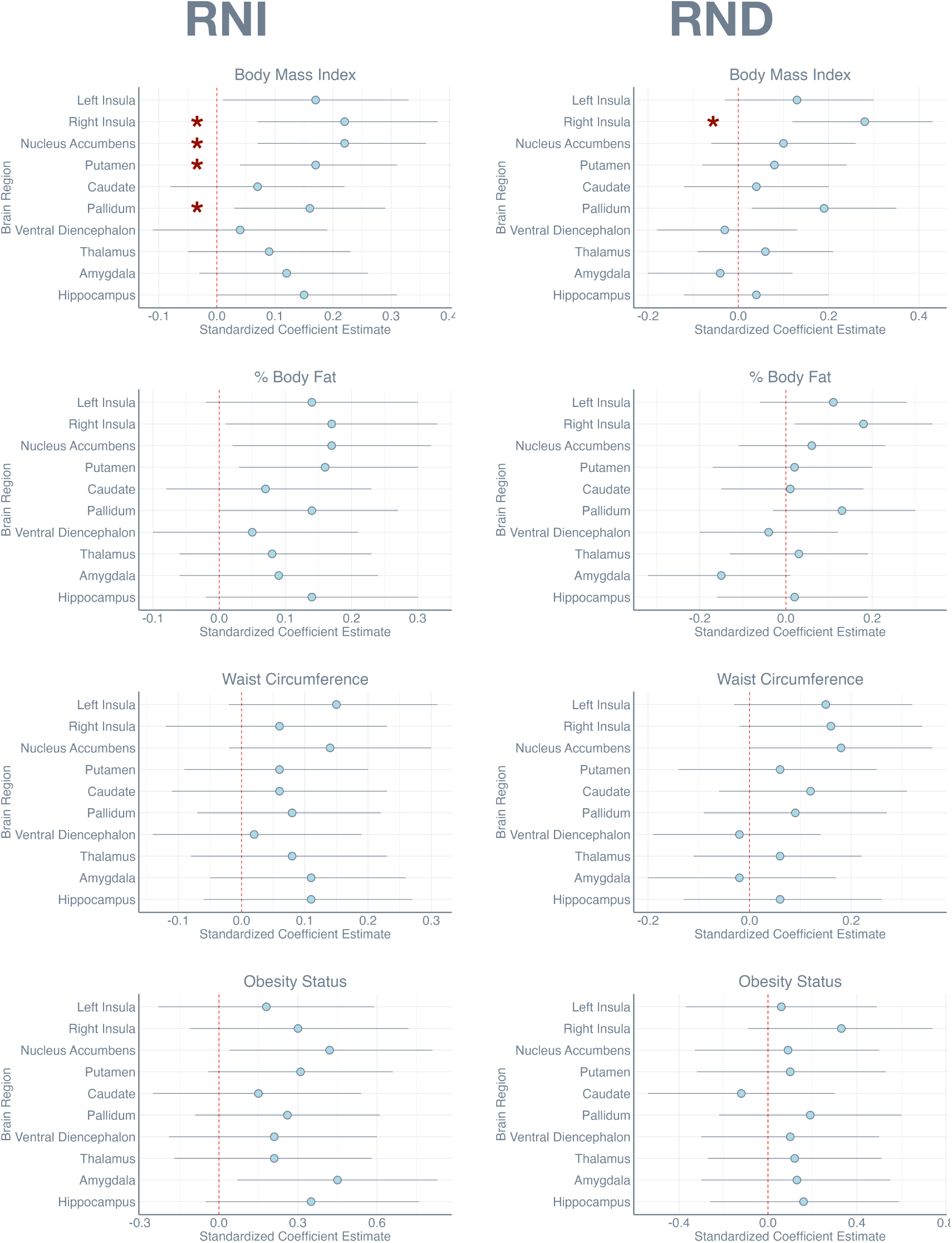
Dot plots show the standardized coefficient estimate *β* for each region of interest for the RNI and RND measures, accompanied by 95%confidence intervals. Red * marks those comparisons that survived False Discovery Rate correction.

**FIGURE 2.**
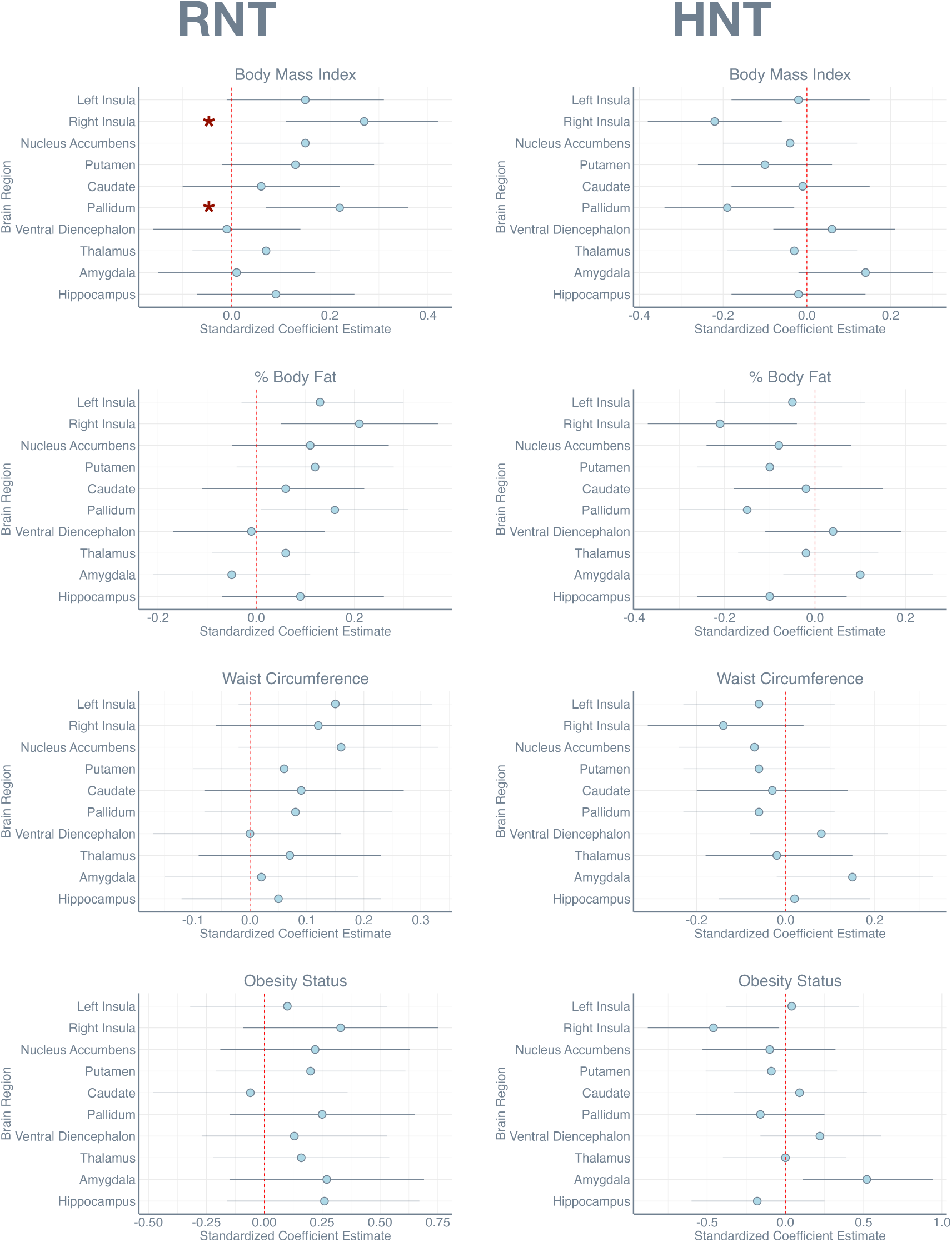
Dot plots show the standardized coefficient estimate *β* for each region of interest for the RNT and HNT measures, accompanied by 95%confidence intervals. Red * marks those comparisons that survived False Discovery Rate correction.

Finally, Figures 3 and 4 show spatially where these effects are strongest for the BMI measure. In a., data are unthresholded, masked for insula, NAcc, putamen, caudate, pallidum, ventral diencephalon, thalamus, amygdala, and hippocampus from the ABCD atlas space. In b., data are thresholded *p* < .01, with the listed regions outlined. Results are consistent with the ROI analysis, showing the strongest and most reliable effects in right insula, in NAcc, and in putamen and pallidum.

**FIGURE 3.**
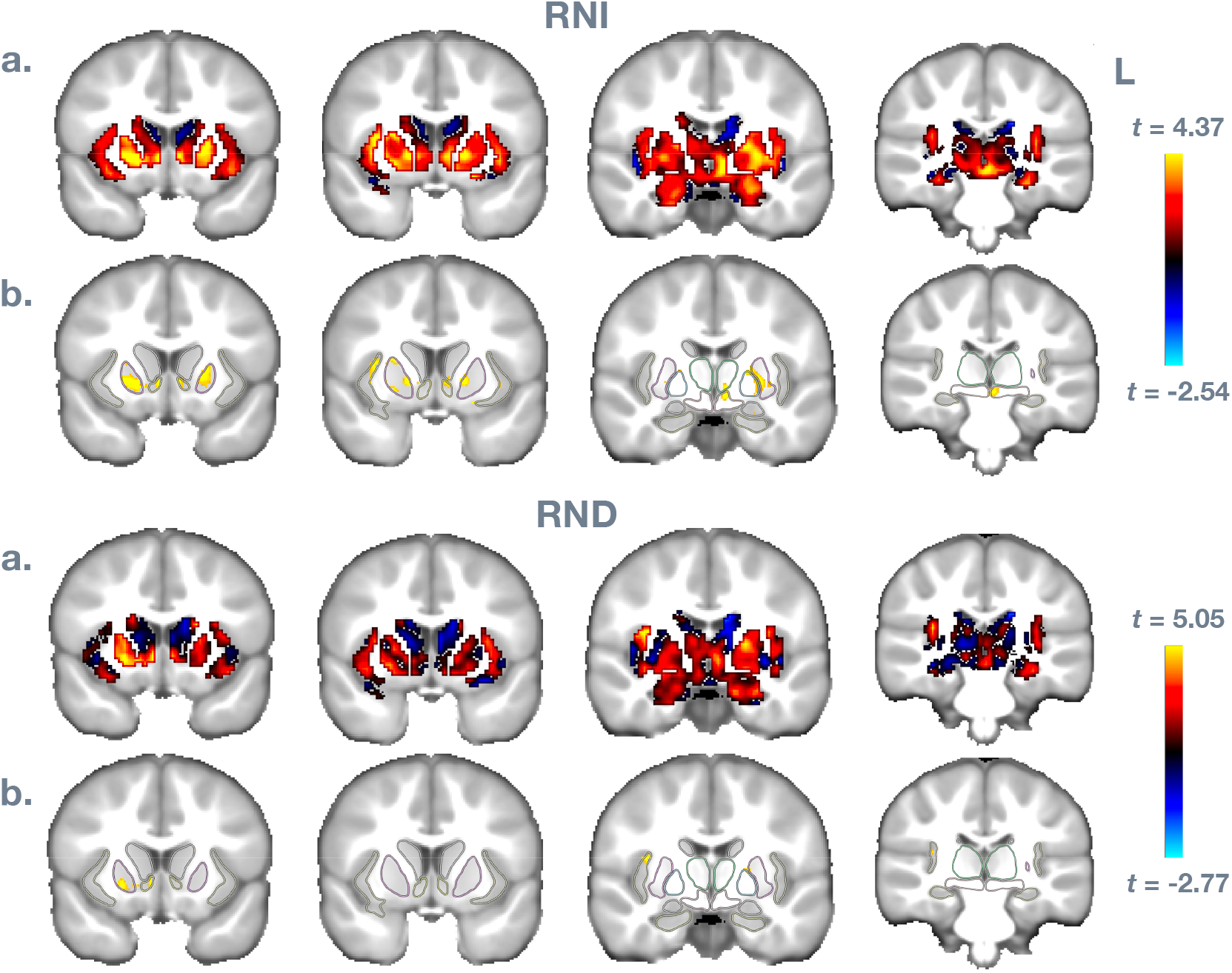
Voxel-wise results for Body Mass Index predicting RNI and RND measures. In a., data are unthresholded, masked for insula, NAcc, putamen, caudate, pallidum, ventral diencephalon, thalamus, amygdala, and hippocampus from the ABCD atlas space. In b., data are thresholded *p* < .01, with the listed regions outlined.

**FIGURE 4.**
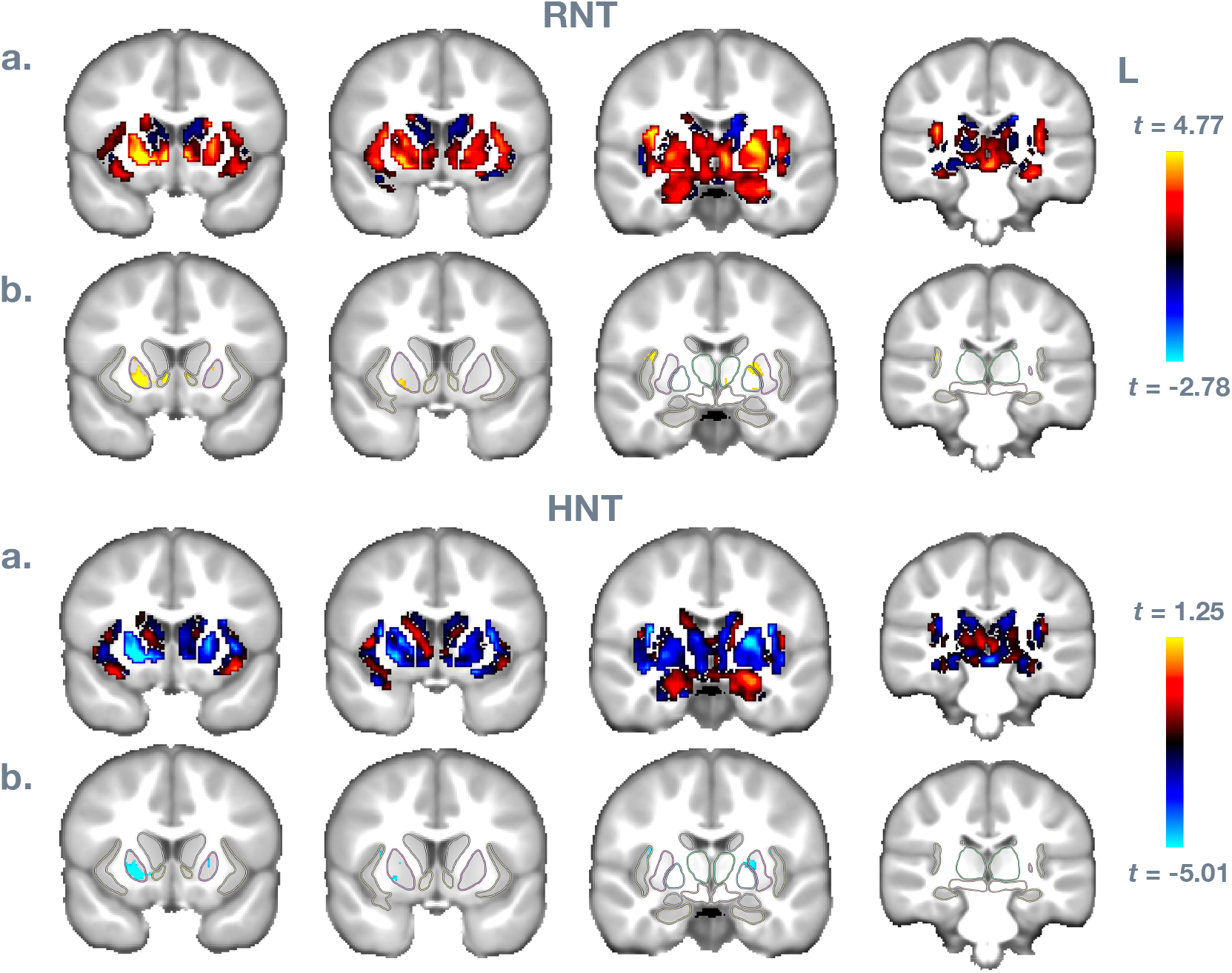
Voxel-wise results for Body Mass Index predicting RNT and HNT measures. In a., data are unthresholded, masked for insula, NAcc, putamen, caudate, pallidum, ventral diencephalon, thalamus, amygdala, and hippocampus from the ABCD atlas space. In b., data are thresholded *p* < .01, with the listed regions outlined.

## 4 DISCUSSION

This study examined how childhood adiposity relates to brain microstructure using RSI metrics in a sample of very young children (ages 4–7 years), including children with and without ADHD. We found that higher BMI was significantly associated with altered brain microstructure, specifically increased RNI diffusion in key subcortical and cortical regions, notably the nucleus accumbens (NAcc) and anterior (predominantly right) insula, as well as pallidum and putamen (associations with RND and RNT were also found for these regions). Unlike previous studies in older children, which emphasized waist circumference as a reliable predictor of brain differences Rapuano et al. (2020 2022), our findings identified BMI as the most sensitive adiposity indicator in early childhood. Our primary hypotheses regarding associations between NAcc and anterior insula cellular density with adiposity were partially supported, as these relationships were observed only for BMI and not for waist circumference, percent body fat, or obesity status. Contrary to expectations, ADHD diagnosis did not moderate these associations; relationships between BMI and brain microstructure were similar across ADHD and typically developing groups, failing to support our hypothesis of greater associations among children with ADHD. We discuss these findings in turn below.

### 4.1 RSI Metrics Reveal Early Associations of Adiposity with Subcortical and Insular Microstructure: Evidence for a Neuroinflammatory Mechanism

Our results replicate and extend the foundational work of Rapuano and colleagues (2020, 2022) and Adise and colleagues (2025), who reported strong associations between waist circumference and BMI, and RNI within subcortical regions, particularly the NAcc, in older children (adolescents in the ABCD study). In our younger sample, the robust association between BMI and RNI in the NAcc and other basal ganglia structures aligns with these previous findings, especially with the study by Adise and colleagues (2025), suggesting that this relationship emerges early in development. However, in contrast to prior work, we did not find significant associations with waist circumference, percent body fat, or categorical obesity status. This discrepancy may reflect developmental differences in sensitivity and measurement reliability across age groups, underscoring BMI’s stability and utility in young populations. It may be the case that measures like waist circumference become more predictive as children get older (Rapuano et al. 2020).

A critical novel contribution of our study is the clear identification of the insula as a significant cortical region associated with obesity-related microstructural differences, which was not highlighted in earlier RSI studies. The insula plays a central role in taste perception and salience processing (Pritchard et al. 1999, Berthoud 2012, Rolls 2016), and functional imaging consistently identifies increased insular responses to palatable food signals among adults and children with obesity (Boutelle et al. 2015, Nakamura and Koike 2021, Avery et al. 2021, Doornweerd et al. 2018, Rapuano et al. 2023). Our findings, demonstrating increased RNI diffusion in the anterior insula, suggest that structural microarchitectural changes may underlie or accompany these functional patterns. RSI separates diffusion signals into biologically informative compartments—restricted, hindered, and free—with the RNI metric specifically isolating isotropic intracellular diffusion. Palmer et al. (2022) and White et al. (2014, 2013b, 2017, 2013a) suggest that RNI-based restricted diffusion is particularly sensitive to cellular density (Yeh et al. 2017, Garic et al. 2021), especially within glial cell populations (Brunsing et al. 2017, Bihan 1995), which may proliferate in response to inflammatory or metabolic signals (Yeh et al. 2017, Garcia-Hernandez et al. 2022).

Indeed, Rapuano et al. (2020, 2022) proposed a model in which poor diet initiates systemic inflammation that crosses the blood–brain barrier and induces glial proliferation and neuroinflammatory responses in regions involved in reward and salience processing. In our study, this mechanism was most evident in the nucleus accumbens (NAcc) and anterior insula, but extended to other subcortical structures including the pallidum and putamen. According to this model, inflammation-driven glial changes may heighten responsivity to food-related cues and disrupt regulatory control, promoting weight gain and reinforcing a feed-forward cycle of maladaptive eating behavior and neural dysregulation. These effects may be particularly pronounced in the anterior insula and NAcc given their dense vascularization, high metabolic demand, and roles in taste perception, interoception, and reward valuation. Our finding that RNI was the most predictive RSI metric reinforces this interpretation. As emphasized by Palmer et al. (2022), RNI reflects isotropic intracellular diffusion and is especially sensitive to changes in glial cell density and morphology—features that are widely recognized as hallmarks of neuroinflammation (Yi et al. 2019, Yang et al. 2013, Turkheimer et al. 2023, Adise et al. 2025). Together, these results support the hypothesis that diet-related neuroinflammatory processes may already be influencing brain structure by early childhood, potentially biasing children toward heightened reactivity to palatable food cues and reduced inhibitory control.

While neuroinflammation is a compelling explanation for increased RNI, non-inflammatory mechanisms may also contribute to our observed diffusion differences. For example, there are at least two possibilities: (1) *Glial proliferation independent of inflammation*—such as astrocytic or microglial expansion in response to energy demands—may also increase intracellular diffusion signals without triggering an immune response (Douglass et al. 2017). (2) *Experience-dependent synaptic plasticity*, whereby frequent engagement with rewarding dietary stimuli strengthens excitatory connections within NAcc and insular networks, could reshape cellular architecture in ways detectable by RSI (Lehmann et al. 2023). These possibilities are not mutually exclusive and may operate in parallel with inflammatory processes. Future multimodal research incorporating metabolic, immune, and functional data is essential to disentangle these mechanisms.

### 4.2 Lack of Moderation by ADHD: Common Neural Pathways in Early Childhood

Contrary to our initial hypothesis, we found no evidence that ADHD diagnosis moderated the relationship between BMI and RNI. Prior research has linked ADHD to altered reward circuitry and increased obesity risk (Cortese and Tessari 2017, Volkow et al. 2013), raising the possibility of shared neurobiological mechanisms. However, our findings suggest that early obesity-related microstructural differences in regions such as the NAcc and insula are comparable between children with ADHD and their typically developing peers. This points to a potentially common neural pathway underlying adiposity in early childhood, regardless of diagnostic status. Notably, we also found no reliable group differences in anthropometric measures, further indicating limited ADHD-related divergence at this developmental stage.

It is possible that neurobiological differences associated with ADHD and obesity become more pronounced later in development (Martins-Silva et al. 2022). Indeed, a meta-analysis by Nigg et al. (2016) reported no clear association between ADHD and obesity in children ages 5– 12, with effect sizes increasing slightly in adulthood (odds ratios rising from 1.17 in childhood to 1.37 in adults). It is also possible that only a subset of individuals with ADHD may be particularly vulnerable to obesity—possibly those with co-occurring internalizing symptoms or other comorbidities (Nigg et al. 2016, Garcia-Hernandez et al. 2022).

While ADHD status did not moderate brain–BMI associations, our study design allowed us to test this hypothesis in a developmentally sensitive way. We included multiple measures of adiposity—BMI, waist circumference, and percent body fat, but found significant effects only for BMI. Although this null finding may reflect weaker predictive utility of alternative measures in this age range, it also reinforces the robustness of the BMI–brain associations observed across both diagnostic groups. Waist circumference and body fat percentage may be more prone to measurement variability in young children due to compliance, hydration, and postural influences (Curzon et al. 2024, Wells and Fewtrell 2006), and may only emerge as stronger predictors in later childhood and adolescence. Nonetheless, the consistency of our findings across ADHD and TD groups supports the idea of early, diagnosis-independent neurobiological markers of obesity risk.

### 4.3 Early Detection and Limitations: The Need for Longitudinal and Multimodal Approaches

Although Rapuano et al. (2020, 2022) leveraged a longitudinal design to demonstrate reciprocal relationships between NAcc microstructure and adiposity over time, our findings show that significant microstructural alterations, particularly in the NAcc and insula, are already observable in a cross-sectional snapshot of children aged 4–7. This suggests that neural signatures associated with obesity risk may emerge earlier than previously documented and could precede overt behavioral or physical manifestations of obesity. Identifying these early neural correlates is critical for informing preventive strategies and understanding the developmental origins of obesity.

However, several limitations constrain the interpretation of these findings. Most notably, our cross-sectional design limits causal inference; longitudinal follow-up is necessary to determine whether early microstructural differences predict future changes in adiposity or reflect early consequences of increased BMI. Additionally, while RNI provides biologically meaningful insights into tissue microstructure—potentially indexing neuroinflammation or glial proliferation, it remains an indirect proxy. Integrating RSI data with metabolic, inflammatory, and functional neuroimaging measures in future multimodal studies will be important for specifying the precise biological mechanisms at play. Finally, although our sample was racially and socioeconomically diverse, replication in larger and independent cohorts is essential to establish the robustness and generalizability of these early neural markers of obesity risk.

### 4.4 Conclusions and Implications for Early Intervention

In summary, our findings indicate that higher BMI in early childhood is associated with significant microstructural changes in reward-related and taste perception/salience processing brain regions, particularly the NAcc and anterior insula. These results strongly support theories linking obesity risk to early neuroinflammatory processes or other cellular alterations detectable via RSI metrics. Importantly, these associations were consistent across ADHD and TD groups, highlighting potential universal neural markers of obesity risk that emerge early in development. Identifying these neural correlates could inform targeted interventions to mitigate obesity risk before it becomes entrenched.

## ACKNOWLEDGEMENTS

We thank the families and children who participated, and continue to participate, in the AHEAD study, as well as staff involved in data collection. Study materials can be accessed by contacting the Corresponding Author at. No part of the study was pre-registered prior to the research being conducted. This research was partially supported by the Intramural Research Program of the National Institute of Biomedical Imaging and Bioengineering in the National Institutes of Health. The views, information or content, and conclusions presented do not necessarily represent the official position or policy of, nor should any official endorsement be inferred on the part of the U.S. government. The research was supported by National Institutes of Health, Grant/Award Numbers: R01MH112588, R01DK119814, R56MH108616.

## AUTHOR CONTRIBUTIONS STATEMENT

A.S.D and P.G. conceptualized and designed the study. A.S.D., M.B., B.B. analyzed the data and wrote the draft manuscript. M.C., M.H., M.C., C.C., and D.G. contributed to data collection and data curation. R. W and M. O. I. contributed to the analysis pipeline. All authors reviewed and commented on the draft manuscript and all authors reviewed and approved the manuscript.

## DECLARATION OF GENERATIVE AI AND AI-ASSISTED TECHNOLOGIES IN THE WRITING PROCESS

During the preparation of this work the authors used ChatGPT in order to improve grammar and flow of content originally-drafted by the authors, or to help with R coding. After using this tool, the authors reviewed and edited the content as needed and take full responsibility for the content of the publication.

