## Supplemental Tables for "Brain Microstructure and Obesity Risk in Early Childhood: Insights from Restriction Spectrum Imaging"

**TABLE 1** Associations of anthropometric measures with RNT

| Brain Region | $\beta$ | SE | $t$ | $p$ | 95% CI Lower | 95% CI Upper | $r_{sp}$ | $FDR_p$ |
|---|---|---|---|---|---|---|---|---|
| **Body Mass Index** | | | | | | | | |
| Left Insula | 0.15 | 0.08 | 1.84 | 0.07 | -0.01 | 0.31 | 0.15 | 0.14 |
| Right Insula | 0.27 | 0.08 | 3.39 | 0.00 | 0.11 | 0.42 | 0.27 | 0.01 |
| Nucleus Accumbens | 0.17 | 0.08 | 2.18 | 0.03 | 0.02 | 0.33 | 0.17 | 0.10 |
| Putamen | 0.14 | 0.08 | 1.83 | 0.07 | -0.01 | 0.30 | 0.15 | 0.14 |
| Caudate | 0.07 | 0.08 | 0.86 | 0.39 | -0.09 | 0.23 | 0.07 | 0.49 |
| Pallidum | 0.21 | 0.08 | 2.71 | 0.01 | 0.06 | 0.36 | 0.21 | 0.04 |
| Ventral Diencephalon | -0.02 | 0.08 | -0.25 | 0.80 | -0.17 | 0.13 | -0.02 | 0.89 |
| Thalamus | 0.09 | 0.08 | 1.23 | 0.22 | -0.06 | 0.24 | 0.10 | 0.37 |
| Amygdala | -0.00 | 0.08 | -0.02 | 0.99 | -0.16 | 0.16 | -0.00 | 0.99 |
| Hippocampus | 0.08 | 0.08 | 0.97 | 0.34 | -0.08 | 0.24 | 0.08 | 0.48 |
| **Waist Circumference** | | | | | | | | |
| Left Insula | 0.16 | 0.09 | 1.83 | 0.07 | -0.01 | 0.33 | 0.15 | 0.29 |
| Right Insula | 0.14 | 0.09 | 1.52 | 0.13 | -0.04 | 0.32 | 0.14 | 0.29 |
| Nucleus Accumbens | 0.18 | 0.09 | 2.03 | 0.04 | 0.00 | 0.36 | 0.18 | 0.29 |
| Putamen | 0.07 | 0.09 | 0.78 | 0.44 | -0.11 | 0.25 | 0.07 | 0.54 |
| Caudate | 0.13 | 0.09 | 1.36 | 0.18 | -0.06 | 0.31 | 0.12 | 0.29 |
| Pallidum | 0.13 | 0.09 | 1.48 | 0.14 | -0.04 | 0.30 | 0.13 | 0.29 |
| Ventral Diencephalon | -0.00 | 0.08 | -0.05 | 0.96 | -0.17 | 0.16 | -0.00 | 0.96 |
| Thalamus | 0.11 | 0.08 | 1.36 | 0.17 | -0.05 | 0.27 | 0.12 | 0.29 |
| Amygdala | 0.06 | 0.09 | 0.70 | 0.49 | -0.12 | 0.25 | 0.06 | 0.54 |
| Hippocampus | 0.11 | 0.10 | 1.14 | 0.26 | -0.08 | 0.30 | 0.11 | 0.37 |
| **% Body Fat** | | | | | | | | |
| Left Insula | 0.12 | 0.09 | 1.45 | 0.15 | -0.04 | 0.29 | 0.12 | 0.37 |
| Right Insula | 0.17 | 0.08 | 2.09 | 0.04 | 0.01 | 0.34 | 0.17 | 0.22 |
| Nucleus Accumbens | 0.12 | 0.08 | 1.45 | 0.15 | -0.04 | 0.29 | 0.12 | 0.37 |
| Putamen | 0.08 | 0.09 | 0.94 | 0.35 | -0.09 | 0.26 | 0.09 | 0.58 |
| Caudate | 0.05 | 0.08 | 0.56 | 0.58 | -0.12 | 0.22 | 0.05 | 0.58 |
| Pallidum | 0.17 | 0.08 | 2.03 | 0.04 | 0.00 | 0.33 | 0.17 | 0.22 |
| Ventral Diencephalon | -0.04 | 0.08 | -0.56 | 0.58 | -0.20 | 0.11 | -0.05 | 0.58 |
| Thalamus | 0.06 | 0.08 | 0.72 | 0.47 | -0.10 | 0.22 | 0.06 | 0.58 |
| Amygdala | -0.10 | 0.09 | -1.13 | 0.26 | -0.27 | 0.07 | -0.10 | 0.52 |
| Hippocampus | 0.05 | 0.09 | 0.60 | 0.55 | -0.12 | 0.22 | 0.05 | 0.58 |
| **Categorized as Obese** | | | | | | | | |
| Left Insula | 0.04 | 0.08 | 0.48 | 0.63 | -0.12 | 0.20 | 0.04 | 0.70 |
| Right Insula | 0.12 | 0.08 | 1.55 | 0.12 | -0.03 | 0.28 | 0.12 | 0.53 |
| Nucleus Accumbens | 0.07 | 0.08 | 0.88 | 0.38 | -0.09 | 0.22 | 0.07 | 0.53 |
| Putamen | 0.09 | 0.08 | 1.12 | 0.26 | -0.07 | 0.24 | 0.09 | 0.53 |
| Caudate | 0.01 | 0.08 | 0.09 | 0.93 | -0.15 | 0.17 | 0.01 | 0.93 |
| Pallidum | 0.11 | 0.08 | 1.43 | 0.15 | -0.04 | 0.26 | 0.12 | 0.53 |
| Ventral Diencephalon | 0.06 | 0.08 | 0.80 | 0.42 | -0.09 | 0.21 | 0.07 | 0.53 |
| Thalamus | 0.08 | 0.07 | 1.11 | 0.27 | -0.06 | 0.23 | 0.09 | 0.53 |
| Amygdala | 0.09 | 0.08 | 1.12 | 0.27 | -0.07 | 0.24 | 0.09 | 0.53 |
| Hippocampus | 0.07 | 0.08 | 0.91 | 0.36 | -0.09 | 0.23 | 0.07 | 0.53 |

**TABLE 2** Associations of anthropometric measures with RNI

| Brain Region | $\beta$ | SE | $t$ | $p$ | 95% CI Lower | 95% CI Upper | $r_{sp}$ | FDR$_p$ |
|---|---|---|---|---|---|---|---|---|
| **Body Mass Index** | | | | | | | | |
| Left Insula | 0.17 | 0.08 | 2.15 | 0.03 | 0.01 | 0.33 | 0.17 | 0.07 |
| Right Insula | 0.22 | 0.08 | 2.80 | 0.01 | 0.07 | 0.38 | 0.22 | 0.03 |
| Nucleus Accumbens | 0.22 | 0.07 | 2.99 | 0.00 | 0.08 | 0.37 | 0.24 | 0.03 |
| Putamen | 0.18 | 0.07 | 2.68 | 0.01 | 0.05 | 0.31 | 0.21 | 0.03 |
| Caudate | 0.09 | 0.08 | 1.22 | 0.23 | -0.06 | 0.25 | 0.10 | 0.28 |
| Pallidum | 0.15 | 0.07 | 2.17 | 0.03 | 0.01 | 0.28 | 0.17 | 0.07 |
| Ventral Diencephalon | 0.03 | 0.08 | 0.40 | 0.69 | -0.12 | 0.18 | 0.03 | 0.69 |
| Thalamus | 0.13 | 0.07 | 1.72 | 0.09 | -0.02 | 0.27 | 0.14 | 0.12 |
| Amygdala | 0.09 | 0.07 | 1.16 | 0.25 | -0.06 | 0.23 | 0.09 | 0.28 |
| Hippocampus | 0.16 | 0.08 | 2.00 | 0.05 | 0.00 | 0.31 | 0.16 | 0.08 |
| **Waist Circumference** | | | | | | | | |
| Left Insula | 0.17 | 0.09 | 1.95 | 0.05 | -0.00 | 0.34 | 0.16 | 0.29 |
| Right Insula | 0.09 | 0.09 | 0.93 | 0.35 | -0.10 | 0.28 | 0.09 | 0.39 |
| Nucleus Accumbens | 0.16 | 0.09 | 1.87 | 0.06 | -0.01 | 0.34 | 0.17 | 0.29 |
| Putamen | 0.10 | 0.08 | 1.28 | 0.20 | -0.05 | 0.25 | 0.11 | 0.29 |
| Caudate | 0.11 | 0.10 | 1.19 | 0.24 | -0.08 | 0.30 | 0.12 | 0.30 |
| Pallidum | 0.10 | 0.08 | 1.29 | 0.20 | -0.05 | 0.25 | 0.11 | 0.29 |
| Ventral Diencephalon | 0.03 | 0.08 | 0.41 | 0.68 | -0.13 | 0.20 | 0.04 | 0.68 |
| Thalamus | 0.13 | 0.08 | 1.63 | 0.10 | -0.03 | 0.29 | 0.14 | 0.29 |
| Amygdala | 0.12 | 0.08 | 1.46 | 0.15 | -0.04 | 0.29 | 0.13 | 0.29 |
| Hippocampus | 0.13 | 0.09 | 1.46 | 0.15 | -0.05 | 0.30 | 0.13 | 0.29 |
| **% Body Fat** | | | | | | | | |
| Left Insula | 0.13 | 0.08 | 1.59 | 0.11 | -0.03 | 0.30 | 0.13 | 0.23 |
| Right Insula | 0.15 | 0.08 | 1.79 | 0.08 | -0.02 | 0.31 | 0.15 | 0.21 |
| Nucleus Accumbens | 0.19 | 0.08 | 2.40 | 0.02 | 0.03 | 0.35 | 0.20 | 0.18 |
| Putamen | 0.16 | 0.07 | 2.10 | 0.04 | 0.01 | 0.30 | 0.18 | 0.19 |
| Caudate | 0.08 | 0.08 | 1.02 | 0.31 | -0.08 | 0.25 | 0.09 | 0.39 |
| Pallidum | 0.12 | 0.07 | 1.73 | 0.09 | -0.02 | 0.27 | 0.15 | 0.21 |
| Ventral Diencephalon | 0.04 | 0.08 | 0.52 | 0.60 | -0.12 | 0.20 | 0.04 | 0.60 |
| Thalamus | 0.10 | 0.08 | 1.28 | 0.20 | -0.05 | 0.25 | 0.11 | 0.29 |
| Amygdala | 0.05 | 0.08 | 0.58 | 0.57 | -0.11 | 0.20 | 0.05 | 0.60 |
| Hippocampus | 0.12 | 0.08 | 1.41 | 0.16 | -0.05 | 0.28 | 0.12 | 0.27 |
| **Categorized as Obese** | | | | | | | | |
| Left Insula | 0.07 | 0.08 | 0.88 | 0.38 | -0.09 | 0.22 | 0.07 | 0.38 |
| Right Insula | 0.11 | 0.08 | 1.45 | 0.15 | -0.04 | 0.27 | 0.12 | 0.21 |
| Nucleus Accumbens | 0.15 | 0.07 | 2.07 | 0.04 | 0.01 | 0.30 | 0.17 | 0.13 |
| Putamen | 0.15 | 0.07 | 2.29 | 0.02 | 0.02 | 0.28 | 0.18 | 0.13 |
| Caudate | 0.08 | 0.08 | 1.07 | 0.28 | -0.07 | 0.23 | 0.09 | 0.32 |
| Pallidum | 0.10 | 0.07 | 1.57 | 0.12 | -0.03 | 0.24 | 0.13 | 0.20 |
| Ventral Diencephalon | 0.08 | 0.07 | 1.06 | 0.29 | -0.07 | 0.23 | 0.09 | 0.32 |
| Thalamus | 0.12 | 0.07 | 1.71 | 0.09 | -0.02 | 0.27 | 0.14 | 0.18 |
| Amygdala | 0.15 | 0.07 | 2.15 | 0.03 | 0.01 | 0.29 | 0.17 | 0.13 |
| Hippocampus | 0.15 | 0.08 | 1.90 | 0.06 | -0.01 | 0.30 | 0.15 | 0.15 |

**TABLE 3** Associations of anthropometric measures with RND

| Brain Region | $\beta$ | SE | $t$ | $p$ | 95% CI Lower | 95% CI Upper | $r_{sp}$ | $FDR_p$ |
|---|---|---|---|---|---|---|---|---|
| **Body Mass Index** | | | | | | | | |
| Left Insula | 0.13 | 0.08 | 1.62 | 0.11 | -0.03 | 0.30 | 0.13 | 0.27 |
| Right Insula | 0.28 | 0.08 | 3.53 | 0.00 | 0.12 | 0.43 | 0.28 | 0.01 |
| Nucleus Accumbens | 0.13 | 0.08 | 1.63 | 0.11 | -0.03 | 0.29 | 0.13 | 0.27 |
| Putamen | 0.08 | 0.08 | 0.99 | 0.32 | -0.08 | 0.24 | 0.08 | 0.62 |
| Caudate | 0.06 | 0.08 | 0.69 | 0.49 | -0.11 | 0.22 | 0.06 | 0.70 |
| Pallidum | 0.19 | 0.08 | 2.34 | 0.02 | 0.03 | 0.34 | 0.19 | 0.10 |
| Ventral Diencephalon | -0.04 | 0.08 | -0.48 | 0.63 | -0.19 | 0.12 | -0.04 | 0.70 |
| Thalamus | 0.07 | 0.08 | 0.89 | 0.37 | -0.08 | 0.22 | 0.07 | 0.62 |
| Amygdala | -0.05 | 0.08 | -0.57 | 0.57 | -0.21 | 0.11 | -0.05 | 0.70 |
| Hippocampus | 0.03 | 0.08 | 0.34 | 0.73 | -0.13 | 0.19 | 0.03 | 0.73 |
| **Waist Circumference** | | | | | | | | |
| Left Insula | 0.15 | 0.09 | 1.68 | 0.09 | -0.03 | 0.32 | 0.14 | 0.31 |
| Right Insula | 0.16 | 0.09 | 1.78 | 0.08 | -0.02 | 0.34 | 0.16 | 0.31 |
| Nucleus Accumbens | 0.18 | 0.09 | 1.94 | 0.05 | -0.00 | 0.35 | 0.17 | 0.31 |
| Putamen | 0.03 | 0.09 | 0.36 | 0.72 | -0.15 | 0.22 | 0.03 | 0.82 |
| Caudate | 0.13 | 0.09 | 1.40 | 0.16 | -0.05 | 0.31 | 0.12 | 0.41 |
| Pallidum | 0.12 | 0.09 | 1.27 | 0.21 | -0.06 | 0.30 | 0.11 | 0.41 |
| Ventral Diencephalon | -0.02 | 0.08 | -0.22 | 0.82 | -0.19 | 0.15 | -0.02 | 0.82 |
| Thalamus | 0.09 | 0.08 | 1.12 | 0.26 | -0.07 | 0.26 | 0.10 | 0.44 |
| Amygdala | 0.02 | 0.09 | 0.25 | 0.80 | -0.16 | 0.21 | 0.02 | 0.82 |
| Hippocampus | 0.09 | 0.10 | 0.88 | 0.38 | -0.11 | 0.29 | 0.08 | 0.55 |
| **% Body Fat** | | | | | | | | |
| Left Insula | 0.11 | 0.09 | 1.33 | 0.18 | -0.06 | 0.28 | 0.11 | 0.46 |
| Right Insula | 0.18 | 0.08 | 2.17 | 0.03 | 0.02 | 0.34 | 0.18 | 0.29 |
| Nucleus Accumbens | 0.08 | 0.09 | 0.88 | 0.38 | -0.09 | 0.24 | 0.07 | 0.63 |
| Putamen | 0.01 | 0.09 | 0.13 | 0.90 | -0.17 | 0.19 | 0.01 | 0.92 |
| Caudate | 0.02 | 0.08 | 0.29 | 0.77 | -0.14 | 0.19 | 0.02 | 0.92 |
| Pallidum | 0.15 | 0.08 | 1.73 | 0.09 | -0.02 | 0.32 | 0.15 | 0.29 |
| Ventral Diencephalon | -0.07 | 0.08 | -0.88 | 0.38 | -0.23 | 0.09 | -0.07 | 0.63 |
| Thalamus | 0.03 | 0.08 | 0.39 | 0.70 | -0.13 | 0.20 | 0.03 | 0.92 |
| Amygdala | -0.16 | 0.08 | -1.88 | 0.06 | -0.33 | 0.01 | -0.16 | 0.29 |
| Hippocampus | 0.01 | 0.09 | 0.11 | 0.92 | -0.16 | 0.18 | 0.01 | 0.92 |
| **Categorized as Obese** | | | | | | | | |
| Left Insula | 0.02 | 0.08 | 0.29 | 0.78 | -0.14 | 0.18 | 0.02 | 0.81 |
| Right Insula | 0.12 | 0.08 | 1.56 | 0.12 | -0.03 | 0.28 | 0.12 | 0.81 |
| Nucleus Accumbens | 0.02 | 0.08 | 0.25 | 0.80 | -0.14 | 0.18 | 0.02 | 0.81 |
| Caudate | -0.02 | 0.08 | -0.24 | 0.81 | -0.18 | 0.14 | -0.02 | 0.81 |
| Pallidum | 0.09 | 0.08 | 1.11 | 0.27 | -0.07 | 0.24 | 0.09 | 0.81 |
| Ventral Diencephalon | 0.05 | 0.08 | 0.65 | 0.51 | -0.10 | 0.20 | 0.05 | 0.81 |
| Thalamus | 0.06 | 0.08 | 0.76 | 0.45 | -0.09 | 0.21 | 0.06 | 0.81 |
| Amygdala | 0.04 | 0.08 | 0.46 | 0.65 | -0.12 | 0.20 | 0.04 | 0.81 |
| Hippocampus | 0.02 | 0.08 | 0.30 | 0.77 | -0.13 | 0.18 | 0.02 | 0.81 |

**TABLE 4** Associations of anthropometric measures with HNT

| Brain Region | $\beta$ | SE | $t$ | $p$ | 95% CI Lower | 95% CI Upper | $r_{sp}$ | $FDR_p$ |
|---|---|---|---|---|---|---|---|---|
| **Body Mass Index** | | | | | | | | |
| Left Insula | -0.02 | 0.08 | -0.18 | 0.85 | -0.18 | 0.15 | -0.01 | 0.91 |
| Right Insula | -0.22 | 0.08 | -2.66 | 0.01 | -0.38 | -0.06 | -0.21 | 0.09 |
| Nucleus Accumbens | -0.04 | 0.08 | -0.45 | 0.66 | -0.20 | 0.13 | -0.04 | 0.91 |
| Putamen | -0.10 | 0.08 | -1.22 | 0.23 | -0.26 | 0.06 | -0.10 | 0.63 |
| Caudate | -0.01 | 0.08 | -0.12 | 0.91 | -0.17 | 0.15 | -0.01 | 0.91 |
| Pallidum | -0.18 | 0.08 | -2.26 | 0.03 | -0.33 | -0.02 | -0.18 | 0.13 |
| Ventral Diencephalon | 0.02 | 0.07 | 0.26 | 0.80 | -0.13 | 0.17 | 0.02 | 0.91 |
| Thalamus | -0.04 | 0.08 | -0.51 | 0.61 | -0.19 | 0.12 | -0.04 | 0.91 |
| Amygdala | 0.09 | 0.08 | 1.15 | 0.25 | -0.07 | 0.25 | 0.09 | 0.63 |
| Hippocampus | 0.01 | 0.08 | 0.15 | 0.88 | -0.15 | 0.18 | 0.01 | 0.91 |
| **Waist Circumference** | | | | | | | | |
| Left Insula | -0.05 | 0.09 | -0.59 | 0.56 | -0.22 | 0.12 | -0.05 | 0.71 |
| Right Insula | -0.15 | 0.09 | -1.70 | 0.09 | -0.32 | 0.02 | -0.14 | 0.46 |
| Nucleus Accumbens | -0.07 | 0.09 | -0.80 | 0.43 | -0.25 | 0.10 | -0.07 | 0.71 |
| Putamen | -0.05 | 0.09 | -0.57 | 0.57 | -0.22 | 0.12 | -0.05 | 0.71 |
| Caudate | -0.02 | 0.09 | -0.28 | 0.78 | -0.19 | 0.15 | -0.02 | 0.78 |
| Pallidum | -0.10 | 0.09 | -1.12 | 0.27 | -0.28 | 0.08 | -0.10 | 0.71 |
| Ventral Diencephalon | 0.06 | 0.08 | 0.71 | 0.48 | -0.10 | 0.21 | 0.06 | 0.71 |
| Thalamus | -0.03 | 0.08 | -0.38 | 0.70 | -0.19 | 0.13 | -0.03 | 0.78 |
| Amygdala | 0.16 | 0.09 | 1.81 | 0.07 | -0.02 | 0.34 | 0.16 | 0.46 |
| Hippocampus | 0.07 | 0.09 | 0.78 | 0.44 | -0.11 | 0.25 | 0.07 | 0.71 |
| **% Body Fat** | | | | | | | | |
| Left Insula | -0.06 | 0.08 | -0.73 | 0.47 | -0.23 | 0.10 | -0.06 | 0.78 |
| Right Insula | -0.21 | 0.08 | -2.53 | 0.01 | -0.38 | -0.05 | -0.21 | 0.12 |
| Nucleus Accumbens | -0.06 | 0.08 | -0.74 | 0.46 | -0.23 | 0.10 | -0.06 | 0.78 |
| Putamen | -0.08 | 0.09 | -0.90 | 0.37 | -0.24 | 0.09 | -0.07 | 0.78 |
| Caudate | -0.01 | 0.08 | -0.06 | 0.95 | -0.17 | 0.16 | -0.01 | 0.95 |
| Pallidum | -0.15 | 0.08 | -1.77 | 0.08 | -0.31 | 0.02 | -0.15 | 0.39 |
| Ventral Diencephalon | 0.02 | 0.08 | 0.20 | 0.84 | -0.14 | 0.17 | 0.02 | 0.94 |
| Thalamus | -0.04 | 0.08 | -0.51 | 0.61 | -0.20 | 0.12 | -0.04 | 0.87 |
| Amygdala | 0.03 | 0.09 | 0.31 | 0.76 | -0.15 | 0.21 | 0.03 | 0.94 |
| Hippocampus | -0.11 | 0.09 | -1.20 | 0.23 | -0.29 | 0.07 | -0.11 | 0.78 |
| **Categorized as Obese** | | | | | | | | |
| Left Insula | 0.02 | 0.08 | 0.20 | 0.84 | -0.14 | 0.18 | 0.02 | 1.00 |
| Right Insula | -0.17 | 0.08 | -2.17 | 0.03 | -0.33 | -0.02 | -0.17 | 0.16 |
| Nucleus Accumbens | -0.01 | 0.08 | -0.09 | 0.93 | -0.17 | 0.15 | -0.01 | 1.00 |
| Putamen | -0.04 | 0.08 | -0.46 | 0.65 | -0.19 | 0.12 | -0.04 | 0.92 |
| Caudate | 0.05 | 0.08 | 0.57 | 0.57 | -0.11 | 0.20 | 0.05 | 0.92 |
| Pallidum | -0.07 | 0.08 | -0.94 | 0.35 | -0.23 | 0.08 | -0.08 | 0.92 |
| Ventral Diencephalon | 0.04 | 0.07 | 0.56 | 0.58 | -0.10 | 0.19 | 0.04 | 0.92 |
| Thalamus | -0.00 | 0.08 | -0.01 | 1.00 | -0.15 | 0.15 | -0.00 | 1.00 |
| Amygdala | 0.17 | 0.08 | 2.17 | 0.03 | 0.02 | 0.33 | 0.17 | 0.16 |
| Hippocampus | -0.06 | 0.08 | -0.70 | 0.48 | -0.22 | 0.10 | -0.06 | 0.92 |
